# Correlative super-resolution analysis of cardiac calcium sparks and their molecular origins in health and disease

**DOI:** 10.1101/2023.02.12.528168

**Authors:** Miriam E. Hurley, Ed White, Thomas M.D. Sheard, Derek Steele, Izzy Jayasinghe

## Abstract

Rapid release of calcium from internal stores via ryanodine receptors (RyRs) is one of the fastest types of cytoplasmic second messenger signalling in excitable cells. In the heart, rapid summation of the elementary events of calcium release, calcium sparks, determine the contraction of the myocardium. Recent advances in super-resolution microscopy methods have revealed complex clustering patterns of RyRs at single-channel resolution. Here, we have experimentally correlated sub-sarcolemmal spontaneous calcium sparks in rat right ventricular myocytes with the underlying nanoscale sub-sarcolemmal RyR patterns resolved with super-resolution DNA-PAINT microscopy. Local functional groupings and clustering patterns of RyR were sampled by the location and the two-dimensional area of each calcium spark to reveal a steep dependence of the integral of a calcium spark on the sum of the local RyRs. Visualisation of recurring spark sites showed evidence of repeated and triggered saltatory activation of multiple local RyR clusters. In similar myocytes taken from rats with induced right ventricular failure, RyR clusters themselves showed a dissipated morphology and fragmented (smaller) clusters. The failing myocytes featured greater heterogeneity in both the spark properties and the relationship between the integral of the calcium spark and the local ensemble of RyRs. Whilst fragmented (smaller) RyR clusters were rarely observed to directly underlying the larger sparks or the recurring spark sites, local interrogation of the channel-to-channel distances confirmed a clear link between the positions of each calcium spark and the tight non-random clustering of the local RyRs in both healthy and failing ventricles.

## Introduction

Clusters of ryanodine receptors (RyRs) form some of the most ubiquitous intracellular calcium (Ca^2+^) signalling nanodomains in excitable cells ^1-5^. The fast Ca^2+^ signals generated can trigger a range of cellular functions including gene transcription ^6^, secretion ^7,8^, drive cellular plasticity, modulate neuronal excitability ^9^ and muscle contraction ^10^. This clustering is particularly critical to the Ca^2+^-induced Ca^2+^ release from the internal stores (primarily the sarcoplasmic-endoplasmic reticulum; SR or ER) into the cytoplasm via type-2 RyR. In striated muscle, elementary Ca^2+^ release events, ‘Ca^2+^ sparks’, summate to give rise to the fast and steep Ca^2+^ transients that activate cellular and, in turn, tissue contraction^11^.

Ca^2+^ sparks have been observed as localised and brief elevations of cytoplasmic Ca^2+^ concentration ([Ca^2+^]_*i*_) and commonly recorded with either confocal ^11,12^, total internal reflection fluorescence (TIRF) ^13^ or types of selective plane illumination microscopies ^14^ for well over three and a half decades. Whilst the size, amplitude and duration of Ca^2+^ sparks vary between cell types, ^2,11^ in striated muscle, they serve as a useful intrinsic bioassay of the performance of the Ca^2+^ handling and the excitation-contraction coupling which it underpins. Deviations in the physical properties of evoked or spontaneous sparks, such as width, duration, latency, shape, integrated [Ca^2+^]_i_, and frequency have often been studied as evidence of dysfunction of the cellular Ca^2+^ handling in cardiac^15-18^ or skeletal^19^ muscle pathologies.

Visualisation of the origins of myocardial Ca^2+^ sparks, the RyRs has advanced considerably in recent years. RyR clusters were first visualised as electron-dense ‘feet’ with scanning and transmission electron microscopy (EM) ^1,20^. Over the past two decades, the spatial organisation of RyR clusters and the molecular components intrinsic to the nanodomains have been described through fluorescence ^21,22^ and tomographic EM ^23-25^. The arrival of super-resolution microscopy and allied functionalities such as target counting, multiplexing and proximity detection have revealed a far more diverse and complex RyR cluster organisation patterns in healthy and failing hearts (see review ^26^). Earlier super-resolution techniques, known by acronyms such as STORM and STED, revealed not only a broader range of RyR cluster sizes, but also the local grouping of sub-clusters (termed ‘super clusters’) which could be functionally coupled by local Ca^2+^ gradients ^17,27^. Some of the second-generation super-resolution microscopy methods that offer resolutions of 15-10 nm, chiefly DNA-point accumulation in nanoscale topography (DNA-PAINT ^28^) and expansion microscopy (ExM ^29^), have allowed the localisation of individual RyR channels within the arrays that constitute these nanodomains. From these studies, we and others have shown that RyRs are arranged in non-uniform patterns, regulated by varying densities of modulatory proteins and may be subjected to heterogeneous post-translational modifications ^30-32^.

A common observation from super-resolution studies of heart pathologies is the fragmented or dissipated morphology of RyR clusters ^17,31-34^. In addition, a diminishing co-localisation of RyRs with their primary triggers ^34,35^, structural and modulatory partner proteins ^32,33^, and the increasing heterogeneity of site-specific RyR-phosphorylation ^31^ have also been reported, coinciding with changes in the Ca^2+^ spark properties. These concurrent shifts in both the structure and function in disease have also renewed interest in the hypothesis that the nano-scale spatial organisation of RyRs must, at least in part, encode the properties of the elementary Ca^2+^ release events. *In silico* simulations have been instrumental in demonstrating that the fragmenting RyR cluster morphologies may impact spark fidelity, amplitude and time course of evoked and spontaneous sparks ^31,36^. They also remain powerful tools for interrogating this structure-function relationship in the context of biochemical modification or multi-molecular partnering that can shift the underpinning determinants of Ca^2+^ sparks (e.g. RyR open probability and luminal [Ca^2+^] of the SR/ER) ^31,37,38^. A significant gap that has remained however is the experimental evidence directly correlating the spark properties observed *in situ* with the true distribution of RyRs.

In this paper, we present a direct correlation of the Ca^2+^ sparks recorded in the sub-sarcolemmal space with the true molecular-scale organisation of peripheral RyRs by leveraging a correlative structure/function imaging protocol that we described recently ^39^. We have overcome the lack of widely-available mammalian models with genetically-encoded reporters RyR or [Ca^2+^]_i 26_ to characterise the shift in the spatial encoding of ventricular Ca^2+^ sparks in right ventricular (RV) failure resulting from pulmonary arterial hypertension (PAH).

## Results

### Pipeline for correlating RyRs underlying peripheral Ca^2+^ sparks

Ca^2+^ sparks are commonly recorded as x-t line-scan confocal kymographs. However, the two-dimensional (2D) spatial patterns of spontaneous Ca^2+^ sparks in the sub-sarcolemmal cytoplasmic space of rat right ventricular myocytes were observable with TIRF microscopy. In TIRF image frames, sparks appear as a transient (∼30-100 ms) and localised event of fluorescence with a full-width-at-half-maximum (FWHM) of 1-6 μm (Fig 1A). The current benchmark for mapping RyR2 channels in cardiomyocytes is DNA-PAINT. With ∼ 10 nm of resolution, DNA-PAINT represents a significant (∼ 25-fold) improvement in optical resolution over standard diffraction-limited methods such as TIRF (Fig 1B). With DNA-PAINT, it is now possible to resolve individual RyRs, observed as punctate labelling densities, that make up each RyR cluster (Fig 1C). In this study, we sought to selectively map the 2D spatial properties of sub-sarcolemmal sparks against the local, 2D organisation of RyRs located within peripheral couplons (schematically illustrated Fig 1D and asterisked region) in healthy and failing RV myocytes using TIRF microscopy.

**Figure 1.**
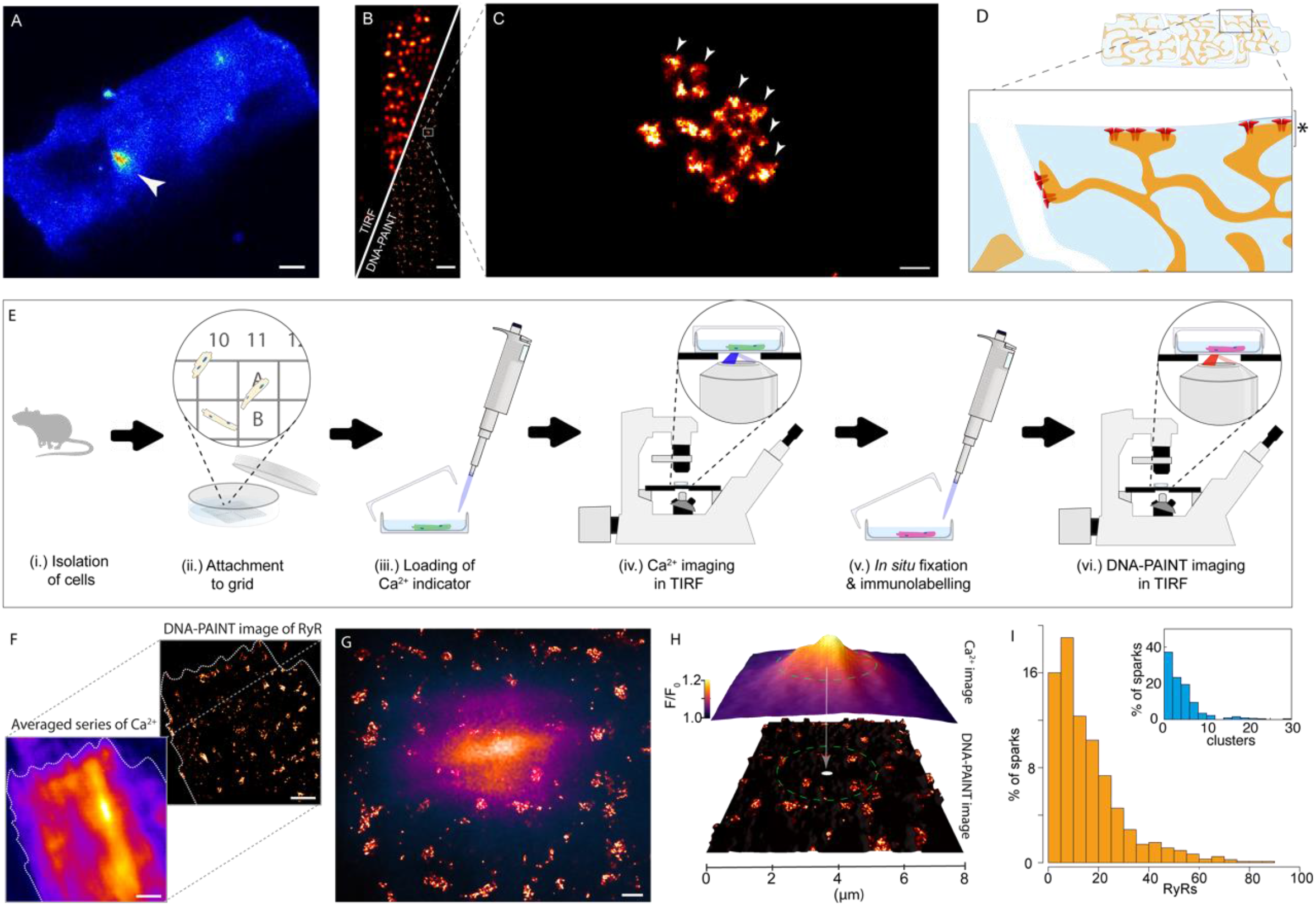
An experimental framework for spatially correlating Ca^2+^ sparks to the underlying RyR channel organisation. A. A TIRF micrograph of a spontaneous Ca^2+^ spark (white arrowhead) in the sub-surface regions of a rat right ventricular myocyte loaded with Fluo-4 AM and immersed in 5 mM [Ca^2+^]_o_. B. A split view of a similar, fixed, rat right ventricular myocyte immuno-stained for RyR2 imaged with standard TIRF microscopy (upper-left) and DNA-PAINT (lower-right), demonstrating the level of resolution improvement in the latter. C. Magnified view of the region indicated by the box in panel B illustrating a single RyR cluster. The individual punctate densities of labelling (arrowheads) represented individual RyR channels; D. A schematic illustration of a ventricular cardiomyocyte where the SR forms junctions with both the surface plasmalemma and the t-tubular invaginations. The magnified view illustrates the predominantly sub-surface RyR clusters observed with the thin TIRF field (depth indicated by asterisk). E. Experimental pipeline developed to allow correlative Ca^2+^ spark TIRF imaging and DNA-PAINT of acutely isolated cardiomyocytes. F. The image alignment involves the upscaling and registration of a time-averaged, two-dimensional version of the Ca^2+^ image series to the density-rendered super-resolution DNA-PAINT image. G. An overlay between a registered Ca^2+^ spark (purple/orange colour-table) and the local super-resolution image of RyR clusters (red hot colour-table). H. Local sampling within the ‘Ca^2+^ spark footprint’ (dashed circle) involved the interrogation of RyR channel and cluster position within a circular window whose diameter was equal to the average FWHM of the spark. I. percentage histograms of the total RyR puncta counts (main panel) and of the total number of unique clusters (inset) within the ‘spark footprint’. Scale bars, A: 5 μm, B: 2 μm, C: 50 nm, F: 1 μm and G: 500 nm.

To this end, we developed a novel experimental pipeline detailed recently ^39^ (Fig 1E). In this protocol, RV cardiomyocytes were isolated freshly from rat hearts, transferred to a glass dish with a gridded bottom, allowing the cells to adhere to the grid. Cytoplasmic fluorescent [Ca^2+^]_i_ indicator Fluo-4 AM was loaded into cells and de-esterified before cells were placed within the TIRF field. Sub-sarcolemmal Ca^2+^ sparks were recorded from cells at specific grid coordinates onto 2D time series. Cells were then fixed *in situ* and labelled with anti-RyR antibodies towards DNA-PAINT imaging and then returned to the TIRF microscope, tracked based on their grid coordinates and DNA-PAINT series recorded. Post-hoc analysis included rescaling and a semi-automated alignment of a time-averaged facsimile of the Ca^2+^ time series against the DNA-PAINT image, primarily using the unique outline of the myocyte to overlay the Ca^2+^ sparks with the underlying RyR map (Fig 1F&G). The coordinates of both the Ca^2+^ sparks and the individual RyR puncta were detected and subsequently registered against each other (using the scaling and alignment vectors established for the whole image). This allowed the RyR channels and clusters within the ‘spark footprint’ (determined by a circular window centred around the spark’s centroid and diameter equal to the spark FWHM) to be counted and sampled on a spark-by-spark basis (Fig 1H). The percentage histograms of the total number of RyR puncta (Fig 1I; main panel) and the number of unique RyR clusters in healthy RV myocytes (i.e. clusters with ≥ 4 RyR puncta; inset) within the footprint of each spark demonstrate that spontaneous Ca^2+^ sparks can arise from broadly varying ensembles of RyR channels (mean of ∼ 18.1 RyRs per spark) and unique clusters (mean ∼ 5.2 clusters per spark; see Supplementary Figure S1-A&C).

### Correlation of RyR organisation with Ca^2+^ sparks in RV failure

To study the structure-function relation of Ca^2+^ sparks in the failing RV, we examined myocytes acutely isolated from the RVs of rats administered with monocrotaline (MCT-RV) and age and sex-matched controls (Ctrl-RV). In the correlative, spark-by-spark analyses, we observed a ∼ 30% increase in the total number of RyRs (mean of ∼ 23.5 RyRs per spark) and a ∼ 60% increase in the number of segmented clusters (mean of ∼ 8.1 clusters per spark; see Supplementary Figure S1-A-D), reflecting the likely recruitment of a greater number of RyRs in the genesis of a spontaneous Ca^2+^ spark. Does this shift in participating RyRs coincide with a *local* change in the overall spark? Spontaneous sparks recorded using TIRF in MCT-RV cells contained on average, ∼38% greater ‘spark mass’ (i.e. the integral of the fluorescence intensity of the spark; see Supplementary Methods section for details) compared to Ctrl-RV (means ∼ 38.4 compared to ∼27.7 respectively). Whilst the FWHM was unchanged, the spark frequency was >90% higher in MCT-RV (see plots and statistics in Supplementary Figure S1E-G).

Close examination of DNA-PAINT images of sub-sarcolemmal RyR from correlative experiments showed a noticeably more dissipated morphology in MCT-RV cells compared to Ctrl-RV (Fig 2-A&B; see arrowheads in panel B). With the use of correlative sparks recorded over ∼20 s, we generated ‘heat maps’ of the local nanoscale RyR morphology (represented by pseudocoloured binary masks of the DNA-PAINT RyR image; see Supplementary Fig S2 for details) to indicate the average estimated spark mass of the locally recorded spontaneous Ca^2+^ sparks in Ctrl-RV and MCT-RV cells (Fig 2-C&D; insets show the greyscale DNA-PAINT images of the shown regions of interest). In the healthy control, the colour-gradations in the spark mass maps were relatively evenly distributed across clusters of similar size whilst smaller, fragmented clusters were sparse. In the failing cells, we observed a large majority of the fragmented (smaller and dispersed) RyR clusters in darker colours, suggesting that they alone are unlikely to produce sparks with greater spark mass. However, regions where higher average spark mass were observed to contain a mix of both fragmented and larger clusters. The scattergrams between the spark mass of each Ca^2+^ spark recorded and the total number of RyRs counted locally revealed a broader degree of scatter in MCT-RV compared to Ctrl-RV (Fig 2E&F). In failing myocytes a considerable number of ‘weaker’ sparks (i.e. spark mass ≤ 1.0) were recorded in regions that contained 20-75 RyRs whilst similar sparks were observed almost exclusively in regions <25 RyR puncta. This noticeable heterogeneity in MCT-RV is further illustrated in the frequency histograms of the total RyR count and the number of unique RyR clusters underneath each spark (Figure 2G&H), represented by the wider distributions of MCT-RV compared to Ctrl-RV. Further, the direct ratio between the recorded spark mass and the locally-counted RyRs showed a lower mean and median for MCT-RV (mean ± SD 1.8 ± 6.7; median 0.53) compared to Ctrl-RV (mean ± SD 2.1 ± 3.1; median 0.53). The increased heterogeneity was represented by an ∼ 2.2-fold higher standard deviation compared to the control (see histogram and violin plot in Fig 2I).

**Figure 2.**
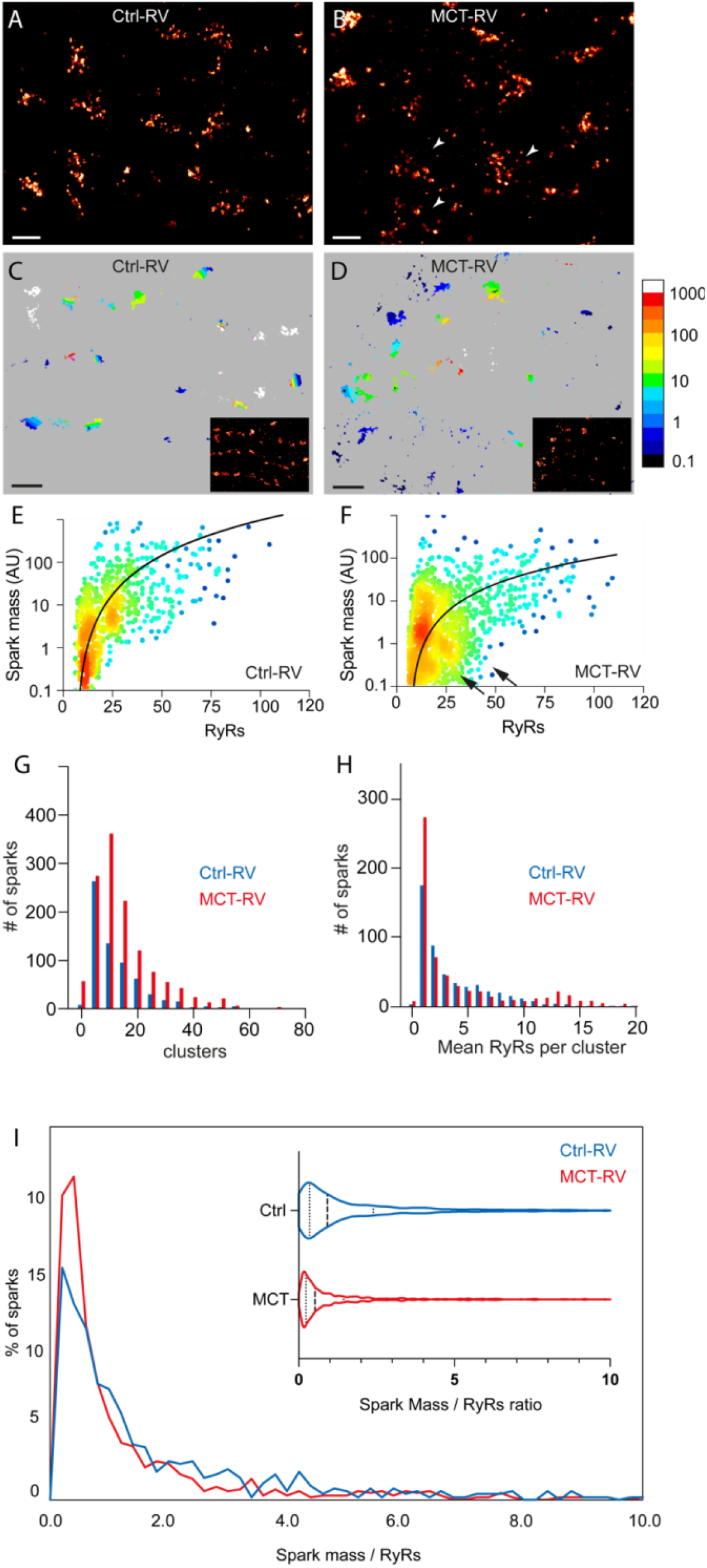
Local interrogation of RyR organisation in Ctrl and MCT-RV cardiomyocytes. DNA-PAINT super-resolution maps of RyR2 labelling in (A) Ctrl and (B) MCT-RV rat cardiomyocytes. The arrowheads indicate the dissipated morphology of RyR clusters in the latter. C&D. show 2D super-resolution maps of RyR, colour-coded for local, average Ca^2+^ spark mass in Ctrl-RV and MCT-RV myocytes respectively. The insets show the original DNA-PAINT images in the corresponding region (larger versions of the insets shown in Supplementary Figure S2); the colour scale represents the average spark mass (in arbitrary units), estimated by xyspark. E&F. Scattergrams of the Ca^2+^ spark mass, plotted on log_10_ scales against the RyRs locally-counted within each ‘spark footprint’ in Ctrl-RV myocytes (n = 1050 sparks; 15 cells; 6 animals) and MCT-RV myocytes (n = 1529 sparks; 15 cells; 6 animals), the latter featuring sparks with smaller spark mass in regions with larger RyR counts. The line-fits are y=0.272x^2^-0.3 and y=0.022x^2^-0.2 respectively. G. Overlaid frequency histograms of the number of RyR clusters consisting of ≥ 5 RyRs within the footprint of sparks recorded in Ctrl-RV (blue; n = 676 sparks) and MCT-RV (red; n = 1326 sparks). H. Overlaid frequency histograms of the mean RyRs detected per cluster within the of sparks recorded in Ctrl-RV and MCT-RV. I. Main panel shows overlays the percentage histograms of the ratio of the estimated spark mass to the locally detected RyRs in Ctrl-RV (blue) and MCT-RV (red). The equivalent violin plot is shown in the insert; medians (0.91 for Ctrl-RV and 0.53 for MCT-RV) in dashed-lines and quartiles in dotted lines. Scale bars: A&B: 200 nm; C&D: 500 nm.

### Relationship between non-random, sub-sarcolemmal Ca^2+^ spark pattern and multi-scale RyR organisation

A novel visualisation unlocked by the correlative approach is the direct overlay of the centroids of all spontaneous Ca^2+^ sparks recorded using TIRF over a window of time onto the 2D super-resolution image of the intricate rows of sub-sarcolemmal RyR clusters (Figure 3A). We investigated the non-random distribution of the spark centroids visually observed across the field of view by performing a 2D quadtree segmentation of domains with higher density of aggregation of spark centroids over a 15 s time window (see Supplementary section 1.9 and Supplementary Fig S3). This segmentation revealed 2D, sub-sarcolemmal regions, typically a few hundred nanometres in width, throughout a cell’s footprint which we classified as ‘recurring spark sites’. Figure 3B&C illustrate the overlays between these recurring spark sites and the super-resolution RyR images in Ctrl-RV and MCT-RV respectively (magnified views shown in panels D&E). Strikingly, we observe that the recurring spark sites rarely overlap with larger RyR clusters directly. Instead, they appear to extend *between* them adjacent clusters in both Ctrl-RV and MCT-RV. There was also no observable, consistent co-location of the fragmented RyRs with the recurring spark sites in MCT-RV. The occupancy of the recurring spark sites between RyR clusters, often within hundreds of nanometres, resembled the shared ‘RyR super cluster domains’ proposed previously as a structural correlate of clusters that are likely to co-activate ^40,41^. It was also conceivable that they aligned with the putative super-clusters of the Ca^2+^ release trigger, L-type Ca^2+^ channels with Ca_v_1.2 observed recently ^42^.

**Figure 3.**
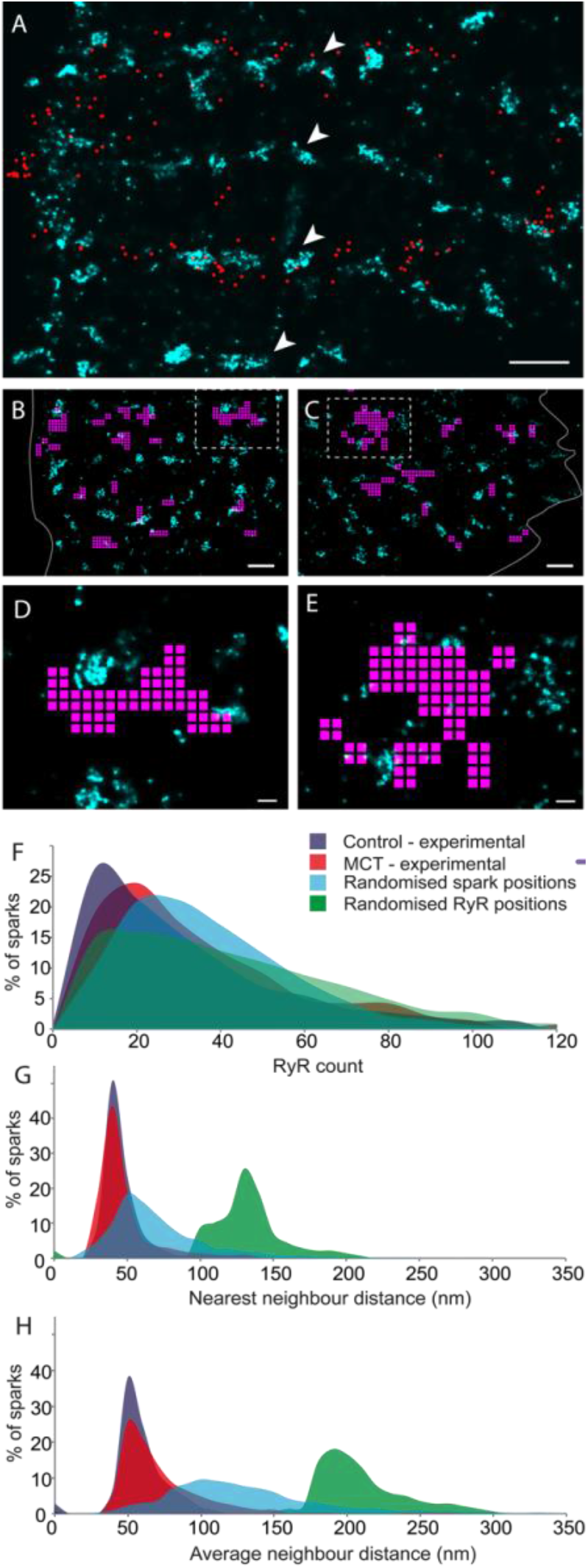
Spatial encoding of the spontaneous Ca^2+^ spark patterns in the RyR organisation. A. An overlay of an RyR2 super-resolution image (cyan) of a Ctrl-RV myocyte and a two-dimensional map of the centroids of all the spontaneous Ca^2+^ sparks (red circles) recorded within a 15 second time window. B&C. Shown, are overlays of the DNA-PAINT super-resolution images of near-surface RyR2 labelling (cyan) and the recurring spark sites (magenta) in Ctrl-RV and MCT-RV myocytes respectively. D&E. Magnified view of the windows indicated by dashed boxes in panels B&C respectively, illustrating that the recurring spark sites typically bridge or tessellate with local groups of RyR cluster rather than overlie them. F. Overlaid percentage histograms of the RyR count underneath the spark footprint in Ctrl-RV (blue) and MCT-RV (red) as well as simulations where the detected spark positions were randomised (cyan), and where RyR positions were randomised. G. The equivalent overlays of the percentage histograms of the mean neighbour distance between RyR puncta underneath each spark, and H the average distance to each of the three nearest neighbours to each RyR underneath spark. Scale bars: A-C 500 nm; D&E: 100 nm.

Both super-resolution and electron tomography image data have confirmed that clustering patterns of RyR channels in healthy cardiomyocytes, albeit not always crystalline, follow a distinct nearest-neighbour distance and the average spacings relative to the surrounding RyRs ^24,30,31^. We sought to investigate whether the spark locations are unique to regions with RyR clustering. To this end, we examined the above variables across experimental datasets (Ctrl-RV and MCT-RV) as well as synthetic images based on control cells where either RyRs or Ca^2+^ sparks were positioned according to random uniform distributions (see Supplementary section 1.10.1 and Supplementary Fig S4). A modest rightward shift and a concurrent broadening in the percentage histograms of the total RyR count (Fig 3F) within each Ca^2+^ spark footprint was observed in both randomised datasets. This reinforces the gain in Ca^2+^ release that is achieved by the clustering of a relatively small number of RyRs compared to randomly and sparsely organised RyRs (see synthetic image in Supplementary Figure S4). In contrast, considerable rightward shifts are observed in both percentage histograms of the nearest neighbour distance of each RyR located under each Ca^2+^ spark, and the average neighbour distance (i.e. average spacing of a given RyR punctum relative to the surrounding RyRs) in clusters detected underneath the spark footprint. This suggests the closeness and uniformity of the intra-cluster organisation of RyRs are intrinsic features critical to producing a spark (Fig 3G&H). Similar to a previous report based on ExM^31^, a modest shift is observed in the rightward tail in the average RyR neighbour distance in MCT-RV (red in Fig 3H) which may reflect that clusters undergo loosening or ‘fraying’ of the RyR positions prior to the fragmentation during RV failure ^32^. Generally, we note the similarity in each of these distributions between Ctrl-RV and MCT-RV datasets in comparison to the randomised datasets. This supports the RyRs remaining within unfragmented clusters as the functionally dominant sub-population of channels driving the genesis of Ca^2+^ spark during disease.

## Discussion

### A novel approach to probe the molecular-scale RyR cluster architecture underlying Ca^2+^ sparks

Presented in this paper is an *experimental* interrogation of the spatial relationship between properties of the elementary events of Ca^2+^ release in primary adult cardiomyocytes and the intrinsic molecular-scale organisation of RyR. Hiess *et. al*. have previously achieved a similar correlation, albeit with TIRF microscopy where GFP-RyR clusters were unresolved ^43^, hence a statistical analysis of the single-channel level RyR organisation was not possible. Our approach was enabled by the development of a correlative imaging protocol^39^ that overcomes the broad unavailability of animal models with endogenous fluorescent reporters of either RyRs and/or [Ca^2+^]_*i*_ that can simultaneously model the cellular and molecular basis of cardiomyopathy. To our knowledge, only DNA-PAINT^30^ and ExM ^32^ currently offers optical resolution sufficient to localise individual RyRs which motivated our choice for correlative DNA-PAINT. The ability to preferentially sample the nanoscale patterns of RyR organisation based on a functional correlate has unlocked a powerful interrogation of the nano/micron-scale structure/function relationship.

Several key conditions and limitations in our approach are however noteworthy. Deviations in the assumed flatness of peripheral RyR clusters, asymmetries and heterogeneities in the 2D spread of Ca^2+^ in the recorded sparks and any acute remodelling RyR clusters within the window of spark imaging (inside 3-4 hours from the point of isolation of cells), albeit minor, are likely sources of error. Whilst Hiess *et. al*. have previously observed with confocal microscopy the movement of some peripheral RyR clusters in quiescent cells maintained under high [Ca^2+^]_*o*_, this fraction was reported to be small ^43^. Although they could reverse this behaviour entirely with exposure to tetracaine, this strategy was incompatible with our objective of observing intrinsically spontaneous sparks. Further, the efficiency of RyR labelling with the use of a widely established antibody (previously^26^ estimated to be at least ∼90%) and the alignment error between the averaged (diffraction-limited) Ca^2+^ image and the super-resolution RyR image can limit the precision of the spark-by-spark sampling of local RyR structure. The key determinants of the latter are the resolution and the signal-to-noise ratio (SNR) of the averaged Ca^2+^ image. Supplementary section 1.10.2 and Supplementary Fig S5 outline a series of simulations by which we demonstrate an alignment error under 100 nm for each iteration given the typical TIRF resolution of ∼ 250 nm and SNR of ∼ 17.0 in our averaged Ca^2+^ image data. For context, an alignment error of 100 nm represents the omission or addition of ∼ 2.5 RyRs in a tightly organised cluster in Ctrl-RV (mean nearest neighbour distance of ∼38.23 nm). Finally, the correlative approach also prevented us from calibrating the pixel values in the Ca^2+^ images against [Ca^2+^]_*i*_ with a Ca^2+^ ionophore due to the requirement of preserving the cellular structure for DNA-PAINT. As a standardisation, all experiments were performed under identical excitation intensities and camera settings which allowed use to quantitatively compare the intensity information Ca^2+^ images between cells and samples.

### Spatial encoding of sparks in the RyR organisation

The non-random dependence of the spark locations on the close RyR clustering particularly reinforces numerous simulations demonstrating how close (and filled) RyR lattice structures promote the ignition of a spark ^31,36,38^. However, similar to previous DNA-PAINT^30^ and ExM^31^ data, our correlative datasets feature a large majority of clusters underneath the spark footprint with looser RyR arrangement (typical neighbour distances > 38 nm) than the fully filled lattices. As confirmed by a recent computational study, this feature is likely to contribute to the considerable variability that we see in the integrated signal of Ca^2+^ sparks ^44^ (e.g. Fig 2E&F) and certainly the broad range of ‘spark mass/RyRs’ ratios that we observe even in the control cells.

The super-positioning of Ca^2+^ spark centroids and the RyR maps (e.g. Fig 3A) provided us an unprecedented view into the spatial encoding of the spark locations across the sarcomerically-organised Ca^2+^ handling machinery. The localisations of the recurring spark sites *between* larger RyR clusters are, to our knowledge, the only *experimental* imaging data supporting the putative ‘triggered saltatory’ recruitment of neighbouring RyR clusters at the genesis of a Ca^2+^ spark ^40^ in both healthy and failing myocytes.

The scattergrams between spark mass and local RyR counts (Fig 2E&F) demonstrate a broad correlation in both failing myocytes and the healthy controls. When the 2D positions of RyRs and sparks were randomised separately, the distribution of the total RyR count within the spark footprint was not entirely annulled (Fig 3F). In fact, it became broader, suggesting that the local RyR ensemble is only a rough determinant of the Ca^2+^ spark. The high degree of scatter in both spark mass scattergrams of Ctrl-RV and MCT-RV (but exacerbated in sparks with higher spark mass in MCT-TV) further reflects how numerically similar pools of RyRs can give rise to Ca^2+^ sparks whose integrated [Ca^2+^]_*i*_ can vary by 2-3 orders of magnitude.

### Heterogeneities in the structure/function relation in RV failure

Increasing structural heterogeneity, including local t-tubule remodelling ^45-47^, local regions of RyR fragmentation ^17,38,48^, diminishing regularity of the expression patterns Ca^2+^ handling proteins over nanometre/micron length-scales^49^, is now understood to be a hallmark of wide-ranging cardiac pathologies. In MCT-RV, we have previously observed evidence of heterogeneous RyR cluster fragmentation ^31^ and the receding sub-domains of RyR-modulators and molecular tethers, junctophilin-2 (JPH2) ^32^ and BIN-1 from some clusters ^18^. In this context, the increasing heterogeneity in the ratio between spark mass and RyR count in MCT-RV (Fig 2I) is the likely result of differently configured *local* Ca^2+^ handling machineries, not limited to the spatial organisation of RyR. Downregulation of the cytoplasmic Ca^2+^ removal mechanisms (the Na^+^/Ca^2+^ exchanger and SR Ca^2+^ ATPase) ^18,50^ along with the overload of SR luminal [Ca^2+^] need to be considered with the 30% increase in the RyR ensemble observed underneath Ca^2+^ sparks. Whilst the individual cluster size of RyR is reduced in MCT-RV as a consequence of the cluster fragmentation or fraying ^31,32^, the modest increases in the RyR-RyR neighbour distance measurements offer some initial clues of the evolving functional coupling in and around the dyads. An enhancement of the functional coupling of the loosely packed RyRs is certainly conceivable with the reduced L-type Ca^2+^ channel expression^50^ and higher SR [Ca^2+^]_*i*_ in MCT-RV myocytes, offering greater driving force during Ca^2+^ release. The super-resolution heatmaps encoding spark mass do not however reveal a clear correlation between fragmented RyRs and regions with higher average spark mass. This observation may however be different if Ca^2+^ leak could be mapped with a more sensitive approach than diffraction-limited TIRF imaging with Fluo-4 Ca^2+^ indicator.

## Materials and Methods

### Microscope setup

All experiments were performed on a Nikon TE2000 (Nikon; Japan) modified to enable total internal reflection fluorescence (TIRF) dual-colour imaging with 488 nm and 671 nm excitations on a ∼ 15 μm x 15 μm illumination field (as detailed previously ^39^). The full list of specifications and settings can be found in the see the supplementary methods section. Emitted light was recorded onto a Zyla 5.5 scientific CMOS camera (sCMOS; Andor, Belfast). Raw image series for both Ca^2+^ imaging and DNA-PAINT were acquired using the open-sources Python Microscopy Environment^51^ (PyME) software.

### Animal models and isolation of ventricular myocytes

Experiments were performed according to the UK Animals (Scientific Procedures) Act of 1986 and with UK Home Office approval and local ethical approval. As a model of right ventricular (RV) failure, adult male Wistar rats aged ∼ 5 weeks were given an intraperitoneal injection of crotaline (Merck, NJ) to induce pulmonary arterial hypertension. Age and sex-matched controls were given an equivalent bolus of saline. At 3-4 weeks, the MCT treated animals were monitored for signs of RV failure were euthanised along with age-matched Ctrl animals. Hearts were dissected acutely and right ventricular myocytes enzymatically isolated following Langendorff perfusion. For a detailed account of the animal model and cell isolation, see the supplementary methods section.

### Ca^2+^ sparks imaging

RV myocytes were loaded with Fluo-4 AM fluorescent Ca^2+^ indicator (ThermoFisher Scientific) and immersed in a Tyrode’s solution containing 5 mM CaCl_2_ at pH 7.4 and adhered to a gridded imaging dish with a #1.5H glass coverslip bottom (Ibidi, USA) for 90 mins before being immersed in fresh Tyrode’s solution for Ca^2+^ spark imaging. The dishes were clamped securely onto the stage of the TIRF microscope system such that the grid was aligned with the straight edges of the camera’s field of view, under brightfield illumination. Cells forming a substantial footprint were illuminated with a 488 nm laser. The local changes in the Fluo-4 fluorescence were recorded at a 100 ms/frame rate with either Tetraspeck microspheres (ThermoFisher Scientific) attached to the imaging grid or the outline of the cell’s contact patch with the grid included within the imaging frame. See detailed protocol in the *supplementary methods* section.

### Cell fixation and immunolabelling for DNA-PAINT imaging of RyR

Immediately following Ca^2+^ imaging, cells were fixed *in situ* with 2% paraformaldehyde (Sigma-Aldrich; w/v in phosphate buffered saline; PBS) for 10 mins at room temperature (RT). Samples were washed and subjected to immunofluorescence labelling with a monoclonal mouse anti-RyR2 IgG (MA3-916; ThermoFisher) primary antibody and an anti-mouse IgG secondary antibody (Jackson Immunoresearch) conjugated to a DNA-PAINT P1 ‘docking’ strand (As designed by Jungmann *et. al*. ^28^) was applied, diluted at 1:100 in incubation solution. See details of the labelling protocol and the production of the secondary antibody and the base sequences of the P1 ‘docking’ strands in the *supplementary methods* section.

### DNA-PAINT imaging and primary processing

Following immunolabelling, samples were washed 3 times in ‘Buffer C’ formulated by Jungmann et al ^28^, and the P1 ‘imager’ strands linked to Atto655 were applied at 1.2 nM in Buffer C. The dish was returned to the microscope stage and the grid coordinates recorded from the Ca^2+^ imaging were used to return the field of view to the corresponding cells imaged previously. For exciting the Atto655 on the imager strands marking the RyR2 targets at the very edge of the cell, the 671 nm laser was focused at a supra-critical angle onto the field of view. Time series of the DNA-PAINT events consisting of 20,000-50,000 frames were acquired at 100 ms/frame integration time. The analysis included the detection of single molecule events and least-squares fitting of a 2D Gaussian to localise their sub-pixel scale centroid. The event positions were then rendered into a 16-bit greyscale TIFF image with a pixel scaling of 5 nm/pixel using an algorithm based on Delaunay triangularisation ^52^.

### Correlative and quantitative image analysis

The full image analysis protocol consisting of the registration of the Ca^2+^ spark data to the DNA-PAINT data and the localisation of Ca^2+^ sparks has been published elsewhere ^39^.

#### Ca^2+^ spark detection

The Ca^2+^ spark localisation tool ImageJ plugin software: ‘xySpark’ ^12^ was used for background estimation, detection and localisation of individual Ca^2+^ sparks. Output of this analysis was a list of x, y and t coordinates of each spark along with their full-width at half maximum (FWHM) estimated by the Gaussian fit, coefficient of determination *R*^*2*^, amplitude estimated as *F*/*F*_*0*_, where *F*_*0*_ was the estimate of the baseline level of the Ca^2+^ indicator fluorescence in a local cytoplasmic region and *F* was the fluorescence intensity value at the peak of the spark. Only sparks with 1.0 *μ*m ≤ FWHM ≤ 6.0 *μ*m **and** an *R*^*2*^ value ≤ 0.5 were filtered and retained for further analysis.

#### RyR puncta localisation

The punctate RyR labelling densities in the rendered images were detected using a centroid detection algorithm available through PyME and describe previously ^30^. The list of coordinates from this analysis were used for the correlative analysis of RyR, analysis of neighbour distances between local RyRs, and counting RyR numbers within cluster and underneath the spark footprints.

#### Image correlation and alignment of discretised Ca^2+^ sparks and RyR puncta

We used an image alignment pipeline code to upscale the Ca^2+^ images and align the coordinates of the Ca^2+^ sparks to the DNA-PAINT RyR image, as described recently ^39^. Briefly, ∼10 consecutive frames from the Ca^2+^ time series were averaged to produce a low-noise, diffraction-limited image of the cell. This image was taken as a reference image of the cell’s relative position against the Tetraspeck microsphere fiduciary markers and the cell boundaries. The code upscaled the low-noise Ca^2+^ image to match the pixel scaling of the DNA-PAINT image. It then required the user to manually align the Ca2+ image against the DNA-PAINT image, using either the cell outlines and/or the Tetraspeck fiduciary markers as guides. The x and y shift coordinates used for this alignment were then used as a starting point to perform an automated fine-alignment through cross-correlation of the images. The shift coordinates determined through the fine alignments were then applied to the Ca^2+^ spark coordinates align them against the maps of RyR puncta. See Hurley, et al. ^39^ for details.

#### Local sampling of RyR organisation using Ca^2+^ spark footprint

RyR centroids and/or segmented RyR clusters located inside a circular window whose diameter was equal to the FWHM of the spark was included in the correlative analysis for each spark. In correlation of the RyR count beneath each spark against ‘spark mass’, the latter was calculated as the product of the spark amplitude (*F*/*F*_*0*_), FWHM^3^ and conversion factor 1.206, and was one of the default output parameters of the xySpark software ^53^. In the analysis of NND and 3ND, only RyR clusters (as defined by each segmented area) that consisted four or more RyRs were considered. The NND value for each spark sampled represented the average of the distance from each RyR centroid to its nearest neighbour. 3ND was the average of the distances from each RyR centroid to each of its three nearest neighbours. The mean of all 3ND estimates for all RyRs found underneath each spark is shown in Figure 3 as a measure of the overall uniformity of RyR arrangement within their clusters, as introduced previously ^26,31^.

#### Cluster segmentation

Using custom-written programs implemented in IDL, a global threshold which encapsulated 80% of the total labelling fraction above background was adopted to generate a mask of the RyR labelled area defining each cluster as per previously published studies ^26,31^.

## Author Contributions and Notes

IJ, EW and DS designed experiments and provided supervision, MEH and TMDS, performed the experiments, DS provided some of the analytical tools, IJ and MEH analysed data, IJ, MEH and DS interpreted the data. IJ and MEH wrote the manuscript.

The authors declare no conflict of interest.

## Acknowledgments

We thank Dr Michael Colman for helpful discussions, Prof Christian Soeller and Dr David Baddeley for advice on implementing the PyME software. The work was funded by the Wellcome Award (207684/Z/17/Z) UKRI award (MR/S03241X/1) made to IJ, and the Leeds Anniversary Research Scholarship awarded to MEH.

## References

1 Sun, X. H. et al. Molecular architecture of membranes involved in excitation-contraction coupling of cardiac muscle. J Cell Biol 129, 659–671, doi:10.1083/jcb.129.3.659 (1995).

2 Ouyang, K. et al. Ca2+ sparks and secretion in dorsal root ganglion neurons. Proc Natl Acad Sci U S A 102, 12259–12264, doi:10.1073/pnas.0408494102 (2005).

3 Pritchard, H. A. T. et al. Nanoscale coupling of junctophilin-2 and ryanodine receptors regulates vascular smooth muscle cell contractility. Proc Natl Acad Sci U S A 116, 21874–21881, doi:10.1073/pnas.1911304116 (2019).

4 Jayasinghe, I. D., Munro, M., Baddeley, D., Launikonis, B. S. & Soeller, C. Observation of the molecular organization of calcium release sites in fast-and slow-twitch skeletal muscle with nanoscale imaging. J R Soc Interface 11, doi:10.1098/rsif.2014.0570 (2014).

5 Johnson, J. D., Kuang, S., Misler, S. & Polonsky, K. S. Ryanodine receptors in human pancreatic beta cells: localization and effects on insulin secretion. FASEB J 18, 878–880, doi:10.1096/fj.03-1280fje (2004).

6 Bracci, L. et al. Ca2+ signaling through ryanodine receptor 1 enhances maturation and activation of human dendritic cells. Journal of Cell Science 120, 2232–2240 %@ 0021-9533, doi:10.1242/jcs.007203 (2007).

7 Shakiryanova, D. et al. Presynaptic ryanodine receptor-activated calmodulin kinase II increases vesicle mobility and potentiates neuropeptide release. J Neurosci 27, 7799–7806, doi:10.1523/JNEUROSCI.1879-07.2007 (2007).

8 Santulli, G. et al. Calcium release channel RyR2 regulates insulin release and glucose homeostasis. J Clin Invest 125, 1968–1978, doi:10.1172/JCI79273 (2015).

9 Bertan, F. et al. Loss of Ryanodine Receptor 2 impairs neuronal activity-dependent remodeling of dendritic spines and triggers compensatory neuronal hyperexcitability. Cell Death Differ 27, 3354–3373, doi:10.1038/s41418-020-0584-2 (2020).

10 Kuo, I. Y. & Ehrlich, B. E. Signaling in muscle contraction. Cold Spring Harb Perspect Biol 7, a006023, doi:10.1101/cshperspect.a006023 (2015).

11 Cheng, H., Lederer, W. J. & Cannell, M. B. Calcium sparks: elementary events underlying excitation-contraction coupling in heart muscle. Science 262, 740–744, doi:10.1126/science.8235594 (1993).

12 Steele, E. M. & Steele, D. S. Automated detection and analysis of Ca(2+) sparks in x-y image stacks using a thresholding algorithm implemented within the open-source image analysis platform ImageJ. Biophysical journal 106, 566–576, doi:10.1016/j.bpj.2013.12.040 (2014).

13 Saeki, T., Suzuki, Y., Yamamura, H., Takeshima, H. & Imaizumi, Y. A junctophilin-caveolin interaction enables efficient coupling between ryanodine receptors and BK<sub>Ca</sub> channels in the Ca<sup>2+</sup> microdomain of vascular smooth muscle. Journal of Biological Chemistry 294, 13093–13105, doi:10.1074/jbc.RA119.008342 (2019).

14 Dvinskikh, L. et al. NM2C. 6 (Optical Society of America).

15 Fowler, E. D. et al. Arrhythmogenic late Ca(2+) sparks in failing heart cells and their control by action potential configuration. Proc Natl Acad Sci U S A 117, 2687–2692, doi:10.1073/pnas.1918649117 (2020).

16 Louch, W. E. et al. Slow Ca(2)(+) sparks de-synchronize Ca(2)(+) release in failing cardiomyocytes: evidence for altered configuration of Ca(2)(+) release units? J Mol Cell Cardiol 58, 41–52, doi:10.1016/j.yjmcc.2013.01.014 (2013).

17 Macquaide, N. et al. Ryanodine receptor cluster fragmentation and redistribution in persistent atrial fibrillation enhance calcium release. Cardiovasc Res 108, 387–398, doi:10.1093/cvr/cvv231 (2015).

18 Fowler, E. D. et al. Beta1-adrenoceptor antagonist, metoprolol attenuates cardiac myocyte Ca(2+) handling dysfunction in rats with pulmonary artery hypertension. J Mol Cell Cardiol 120, 74–83, doi:10.1016/j.yjmcc.2018.05.015 (2018).

19 Wang, X. et al. Uncontrolled calcium sparks act as a dystrophic signal for mammalian skeletal muscle. Nat Cell Biol 7, 525–530, doi:10.1038/ncb1254 (2005).

20 Franzini-Armstrong, C., Protasi, F. & Ramesh, V. Shape, size, and distribution of Ca(2+) release units and couplons in skeletal and cardiac muscles. Biophys J 77, 1528–1539, doi:10.1016/S0006-3495(99)77000-1 (1999).

21 Soeller, C., Crossman, D., Gilbert, R. & Cannell, M. B. Analysis of ryanodine receptor clusters in rat and human cardiac myocytes. Proc Natl Acad Sci U S A 104, 14958–14963, doi:10.1073/pnas.0703016104 (2007).

22 Jayasinghe, I. D., Cannell, M. B. & Soeller, C. Organization of ryanodine receptors, transverse tubules, and sodium-calcium exchanger in rat myocytes. Biophys J 97, 2664–2673, doi:10.1016/j.bpj.2009.08.036 (2009).

23 Hayashi, T. et al. Three-dimensional electron microscopy reveals new details of membrane systems for Ca2+ signaling in the heart. J Cell Sci 122, 1005–1013, doi:10.1242/jcs.028175 (2009).

24 Asghari, P. et al. Cardiac ryanodine receptor distribution is dynamic and changed by auxiliary proteins and post-translational modification. Elife 9, doi:10.7554/eLife.51602 (2020).

25 Colman, M. A., Pinali, C., Trafford, A. W., Zhang, H. & Kitmitto, A. A computational model of spatio-temporal cardiac intracellular calcium handling with realistic structure and spatial flux distribution from sarcoplasmic reticulum and t-tubule reconstructions. PLoS Comput Biol 13, e1005714, doi:10.1371/journal.pcbi.1005714 (2017).

26 Jayasinghe, I. et al. Shining New Light on the Structural Determinants of Cardiac Couplon Function: Insights From Ten Years of Nanoscale Microscopy. Front Physiol 9, 1472, doi:10.3389/fphys.2018.01472 (2018).

27 Baddeley, D., Jayasinghe, I. D., Cremer, C., Cannell, M. B. & Soeller, C. Light-induced dark states of organic fluochromes enable 30 nm resolution imaging in standard media. Biophys J 96, L22–24, doi:10.1016/j.bpj.2008.11.002 (2009).

28 Jungmann, R. et al. Multiplexed 3D cellular super-resolution imaging with DNA-PAINT and Exchange-PAINT. Nat Methods 11, 313–318, doi:10.1038/nmeth.2835 (2014).

29 Tillberg, P. W. et al. Protein-retention expansion microscopy of cells and tissues labeled using standard fluorescent proteins and antibodies. Nat Biotechnol 34, 987–992, doi:10.1038/nbt.3625 (2016).

30 Jayasinghe, I. et al. True Molecular Scale Visualization of Variable Clustering Properties of Ryanodine Receptors. Cell Rep 22, 557–567, doi:10.1016/j.celrep.2017.12.045 (2018).

31 Sheard, T. M. D. et al. Three-Dimensional and Chemical Mapping of Intracellular Signaling Nanodomains in Health and Disease with Enhanced Expansion Microscopy. ACS Nano 13, 2143–2157, doi:10.1021/acsnano.8b08742 (2019).

32 Sheard, T. M. D. et al. Three-dimensional visualization of the cardiac ryanodine receptor clusters and the molecular-scale fraying of dyads. Philos Trans R Soc Lond B Biol Sci 377, 20210316, doi:10.1098/rstb.2021.0316 (2022).

33 Hou, Y. et al. Nanoscale Organisation of Ryanodine Receptors and Junctophilin-2 in the Failing Human Heart. Front Physiol 12, 724372, doi:10.3389/fphys.2021.724372 (2021).

34 Lipsett, D. B. et al. Cardiomyocyte substructure reverts to an immature phenotype during heart failure. J Physiol 597, 1833–1853, doi:10.1113/JP277273 (2019).

35 Song, L. S. et al. Orphaned ryanodine receptors in the failing heart. Proc Natl Acad Sci U S A 103, 4305–4310, doi:10.1073/pnas.0509324103 (2006).

36 Walker, M. A. et al. On the Adjacency Matrix of RyR2 Cluster Structures. PLoS Comput Biol 11, e1004521, doi:10.1371/journal.pcbi.1004521 (2015).

37 Hernandez Mesa, M., van den Brink, J., Louch, W. E., McCabe, K. J. & Rangamani, P. Nanoscale organization of ryanodine receptor distribution and phosphorylation pattern determines the dynamics of calcium sparks. PLoS Comput Biol 18, e1010126, doi:10.1371/journal.pcbi.1010126 (2022).

38 Shen, X. et al. Prolonged beta-adrenergic stimulation disperses ryanodine receptor clusters in cardiomyocytes and has implications for heart failure. Elife 11, doi:10.7554/eLife.77725 (2022).

39 Hurley, M. E. et al. A correlative super-resolution protocol to visualise structural underpinnings of fast second-messenger signalling in primary cell types. Methods 193, 27–37, doi:10.1016/j.ymeth.2020.10.005 (2021).

40 Baddeley, D. et al. Optical single-channel resolution imaging of the ryanodine receptor distribution in rat cardiac myocytes. Proc Natl Acad Sci U S A 106, 22275–22280, doi:10.1073/pnas.0908971106 (2009).

41 Hou, Y., Jayasinghe, I., Crossman, D. J., Baddeley, D. & Soeller, C. Nanoscale analysis of ryanodine receptor clusters in dyadic couplings of rat cardiac myocytes. J Mol Cell Cardiol 80, 45–55, doi:10.1016/j.yjmcc.2014.12.013 (2015).

42 Ito, D. W. et al. beta-adrenergic-mediated dynamic augmentation of sarcolemmal CaV 1.2 clustering and co-operativity in ventricular myocytes. J Physiol 597, 2139–2162, doi:10.1113/JP277283 (2019).

43 Hiess, F. et al. Dynamic and Irregular Distribution of RyR2 Clusters in the Periphery of Live Ventricular Myocytes. Biophys J 114, 343–354, doi:10.1016/j.bpj.2017.11.026 (2018).

44 Cosi, F. G. et al. Multiscale Modeling of Dyadic Structure-Function Relation in Ventricular Cardiac Myocytes. Biophys J 117, 2409–2419, doi:10.1016/j.bpj.2019.09.023 (2019).

45 Biesmans, L. et al. Subcellular heterogeneity of ryanodine receptor properties in ventricular myocytes with low T-tubule density. PLoS One 6, e25100, doi:10.1371/journal.pone.0025100 (2011).

46 Perera, T. et al. Serial block face scanning electron microscopy reveals region-dependent remodelling of transverse tubules post-myocardial infarction. Philos Trans R Soc Lond B Biol Sci 377, 20210331, doi:10.1098/rstb.2021.0331 (2022).

47 Li, H. et al. Cardiac Resynchronization Therapy Reduces Subcellular Heterogeneity of Ryanodine Receptors, T-Tubules, and Ca2+ Sparks Produced by Dyssynchronous Heart Failure. Circ Heart Fail 8, 1105–1114, doi:10.1161/CIRCHEARTFAILURE.115.002352 (2015).

48 Kolstad, T. R. et al. Ryanodine receptor dispersion disrupts Ca(2+) release in failing cardiac myocytes. Elife 7, doi:10.7554/eLife.39427 (2018).

49 Holmes, M. et al. Increased SERCA2a sub-cellular heterogeneity in right-ventricular heart failure inhibits excitation-contraction coupling and modulates arrhythmogenic dynamics. Philos Trans R Soc Lond B Biol Sci 377, 20210317, doi:10.1098/rstb.2021.0317 (2022).

50 Xie, Y. P. et al. Sildenafil prevents and reverses transverse-tubule remodeling and Ca(2+) handling dysfunction in right ventricle failure induced by pulmonary artery hypertension. Hypertension 59, 355–362, doi:10.1161/HYPERTENSIONAHA.111.180968 (2012).

51 Marin, Z. et al. PYMEVisualize: an open-source tool for exploring 3D super-resolution data. Nat Methods 18, 582–584, doi:10.1038/s41592-021-01165-9 (2021).

52 Baddeley, D., Cannell, M. B. & Soeller, C. Visualization of localization microscopy data. Microsc Microanal 16, 64–72, doi:10.1017/S143192760999122X (2010).

53 Hollingworth, S., Peet, J., Chandler, W. K. & Baylor, S. M. Calcium sparks in intact skeletal muscle fibers of the frog. J Gen Physiol 118, 653–678, doi:10.1085/jgp.118.6.653 (2001).

