## Supplementary information for "Correlative super-resolution analysis of cardiac calcium sparks and their molecular origins in health and disease"

### Supporting information

#### 1. Supplementary methods

##### 1.1. Animal models and isolation of ventricular myocytes

All experiments were performed according to the UK Animals (Scientific Procedures) Act of 1986 with approval of the UK Home Office. Adult male Wistar rats, aged ~ 5 weeks (200 g) were given an intraperitoneal injection of crotaline (MCT, Sigma-Aldrich) at a dose of 60 mg per kg of body weight to induce pulmonary arterial hypertension as detailed previously (1). Between days 21 and 28, the MCT administered animals were monitored for signs of right ventricular failure (weight loss greater than 10 g in 24 hrs or over two consecutive days, cold extremities, piloerection, lethargy and dyspnoea) and euthanized for experiments. Concurrent, age-matched control (Ctrl) animals were given an equivalent volume of saline without crotaline in the intraperitoneal injection and euthanized on the same days as MCT animals.

Animals were concussed with a blow to the head and euthanized by cervical dislocation. Hearts were briefly excised, cannulated at the aorta onto a flow-controlled Langendorff perfusion system (detailed previously in the supplementary section of Jayasinghe & Clowsley et al. (2). Briefly, the whole heart was cannulated at the aorta. To clear the coronary circulation, isolation solution with added 0.75 mM $CaCl_2$ was perfused at 7 mL/min. Isolation solution at pH 7.4 constituted of (in mM) 130 NaCl, 1.4 $MgCl_2$, 5.4 KCl, 0.4 $NaH_2PO_4$, 5 HEPES, 10 Glucose, 20 Taurine, 10 Creatine. Once cleared, isolation solution with added 0.1 mM EGTA (Sigma-Aldrich, USA) was perfused for 4 minutes. Enzymatic digestion of the whole heart followed. Isolation solution was perfused for 7 minutes with 1 mg/mL Collagenase Type II (Worthington Biochemicals, USA) and 1.8 mg/mL protease (Sigma-Aldrich, USA). Atria and vasculature were dissected away.

The right ventricle was dissected and placed in a conical flask with a solution identical to that used in the perfused enzymatic digestion. Using a rotary shaker, for 5 minutes, the conical flask was shaken within a 37°C water bath prior to the filtering of enzyme solution through a 200 µm nylon mesh gauze. Collected cardiomyocytes were centrifuged (50 rpm; 45 s), supernatant removed, and cardiomyocytes resuspended within 750 µM $CaCl_2$ containing isolation solution. Remaining tissue was shaken again for 5 min with the above steps repeated until no cell-rich tissue remained.

##### 1.2. Cell preparation for calcium sparks imaging

Prior to live-cell $Ca^{2+}$ imaging, cells were resuspended in a Tyrode's solution containing (in mM) 140 NaCl, 4 $MgCl_2$, 1 KCl, 10 HEPES, 10 Glucose, 5 $CaCl_2$ at pH 7.4. Cells were loaded with 5 µM Fluo-4 AM (ThermoFisher Scientific, UK) for 15 minutes at room temperature (RT). Cells were washed twice with Tyrode's solution to remove excess $Ca^{2+}$ indicator dye prior to a 30-min de-esterification stage at 4°C (3). Following a third wash, cells were transferred onto a 11.9 µg/mL laminin (ThermoFisher Scientific, UK) coated 500 µm square gridded imaging dish with a #1.5H glass coverslip (Ibidi, USA) and incubated at 37°C for 90 mins.

##### 1.3. Microscope setup

All experiments were performed on a modified Nikon TE2000 (Nikon; Japan) modified to enable total internal reflection fluorescence (TIRF) imaging capability (as detailed previously (4)). For calcium spark imaging, a 488 nm Cobolt Jive DPSS 200 mW laser (Cobolt AB, Solna) was used. For DNA-PAINT, a 642 nm Cobolt DPSS 100 mW laser was used. The lasers were focused onto the glass bottom of the dish at a supra-critical angle using a Nikon 1.49NA TIRF objective lens to achieve a field of illumination of ~ 15 µm x 15 µm. A multiband ZT375/488/543/633rpc (Chroma) dichroic mirror and a ZET488/640m dual band emission filter (Chroma). Emitted light was recorded onto a Zyla 5.5 USB scientific CMOS camera (sCMOS; Andor, Belfast).

Image acquisition was done on a Lenovo ThinkStation workstation using an Intel i7 quad-core processor, 32 Gb of DDR3 memory and a 2 Tb solid-state drive running the open-sources Python Microscopy Environment (PyME) software (5) (freely-available for download via www.python-microscopy.org) in a Windows 7 Professional operating system. The PyME 'Acquire' image acquisition interface included a live display of the camera's acquisition at a customizable framerate.

**1.4. Calcium sparks imaging**

The imaging dishes were clamped securely onto the stage of the TIRF microscope system such that the grid was aligned with the straight edges of the camera's field of view, under brightfield illumination. Cells forming a substantial footprint were illuminated with a 488 nm laser at a power of ~$1\times10^5$ W/m$^2$. The local changes in the Fluo-4 fluorescence were recorded at a 100 ms/frame frame rate onto a 12-bit HDF time series image file (.h5 format for PyME). In ideal cases, the calcium time series also captured the TetraSpeck microspheres attached to the coverslip adjacent to the cells. Nevertheless, it was ensured that the image series included a clear view of the outline of the cell's contact patch with the coverslip, used later for aligning the calcium sparks images with the DNA-PAINT image data.

**1.5. Sample processing for correlative DNA-PAINT**

Immediately following calcium imaging, cells were fixed *in situ* with 2% paraformaldehyde (Sigma-Aldrich; w/v in phosphate buffered saline; PBS) for 10 mins at room temperature (RT). Fixative was removed with PBS over 4× 10-min washes at RT and the cells were briefly permeabilized at RT with 0.1% (v/v) Triton X-100 for 10 min. This was followed by the application of a PBS based blocking buffer, for 60 minutes at RT, which contained 10% normal goat serum. The monoclonal mouse anti-RyR2 IgG (MA3-916; Thermo Fisher) primary antibody was diluted at 1:200 and applied to the cell samples in incubation buffer containing (in w/v or v/v) 0.05% $NaN_3$; 2% BSA; 2% NGS; 0.05% Triton X-100 dissolved in PBS. An anti-mouse IgG secondary antibody (Jackson Immunoresearch) conjugated to a DNA-PAINT P1 'docking' strand (sequence: C6 dT-amine - 5' TTA TAC ATC TA 3'; custom made via Integrated DNA Technologies (USA); as designed by Jungmann et al (6)) was applied, again diluted at 1:100 in incubation solution. Cells were incubated in primary and secondary antibodies overnight at 4°C and for 2 hrs at room temperature (RT) respectively.

**1.6. Conjugation and standardizing of DNA-PAINT antibody-oligo markers**

Both the P1 'docking' (stated above) and the complementary 'imager' strands were commercially synthesised and HPLC purified by Eurofins Ltd (Luxembourg). We opted for a direct lightening-link thiolation which links a 5' C6 amine of the docking strands and cysteines of the antibody. In addition, the conjugated docking strands contained a 3' fluorophore which has the advantage that the conjugated antibody markers carry a fluorochrome, Cy5, allowing us to confirm positive staining. This fluorophore does not interfere with later DNA-PAINT imaging since generally the 3'-fluorophore was selected spectrally distinct from the imager fluorophore. The docking strands were conjugated to a goat anti-mouse IgG secondary antibody (affinity purified, azide-free form from Jackson Immunoresearch, PA) using a Thunder-Link® kit (Innova Biosciences, Cambridge). The conjugated antibody was purified from unconjugated docking strands by use of a ThunderLink® kit-based precipitation step and centrifugation at 13,000g. The relative concentration of the antibody and the docking strand (by the 3'-fluorophore) in the conjugate were determined with the use of a NanoDrop 2000 spectrophotometer (Thermo Scientific). Only antibody samples with an oligo: antibody conjugation ratio ≥ 1:1 were used for sample labelling.

##### 1.7. DNA-PAINT imaging and live processing

Following immunolabelling, samples were washed 3 times in 'Buffer C' (PBS with final 500 mM NaCl; pH 8.0) designed previously by Jungmann et al (6) and the P1 'imager' strands (sequence: 5'CTA GAT GTA T3'-Atto655) were applied in Buffer C at a 1.2 nM oligo concentration. The dish was returned to the microscope stage and the grid coordinates recorded from the calcium imaging were used to return the field of view to the corresponding cells imaged previously. Using the 'live-view' of the field of view under brightfield illumination, the gridlines were aligned with the edges of the camera image before clamping down the dish onto the stage.

The contact patch between the cell and the coverslip were moved into the centre of the laser spot. For exciting the Atto-655 on the imager strands marking the RyR2 targets at the very edge of the cell, the 671 nm laser was focused at a supra-critical angle onto the field of view at a power of ~$1 – 5 \times 10^6$ W/m$^2$. Time series of the DNA-PAINT events were acquired at 100 ms/frame integration time using using PyME, and saved in either HDF or TIFF image format, typically containing ~ 20,000-50,000 frames.

The frame data of DNA-PAINT single molecule events were analysed in real-time (during acquisition) using PyME. The analysis included the detection of single molecule events and least-squares fitting of a 2D Gaussian to localise their sub-pixel scale centroid. The point data of the localised marker positions from a sub-series of 10,000 frames were selected for further analysis and events lasting more than one frame (tracked in consecutive frames to be within the localisation precision) were coalesced to minimise localisation errors. The event positions were then rendered into a 16-bit greyscale TIFF image with a pixel scaling of 5 nm/pixel using an algorithm based on Delaunay triangularisation (7) such that the pixel intensity was linearly proportional to the local density of localised markers.

##### 1.8. Spark mass heatmaps

Following the correlative image-rescaling, alignment and spark detection, two exemplar datasets consisting of at ~ 20 s of $Ca^{2+}$ spark recordings were selected from each of Ctrl-RV and MCT-RV. The greyscale-rendered DNA-PAINT images of RyR (Suppl Fig S2 A&E respectively) were converted into binary masks by applying a global threshold which encapsulated 80% of the total labelling fraction above background (Suppl Fig S2 B&F respectively). The registered coordinates (post-rescaling and alignment) of the centroid of each $Ca^{2+}$ spark were mapped onto a pixel on a separate, floating-point array. The pixel was assigned a value equal to the spark mass estimated by the xyspark software (freely downloadable from https://xyspark.leeds.ac.uk/) for the original event. Each of these arrays were then convolved with a 2D Gaussian function with a FWHM of 1 µm and re-coloured using the Rainbow lookup table in ImageJ v2.3.0 to obtain the representative spark-mass maps (Suppl Fig S2 C&G respectively). Multiplying these by the corresponding RyR binary mask produced the RyR heatmaps colour-coded for average spark mass- (Suppl Fig S2 D&H).

##### 1.9. Recurring spark sites

Similar to the approach for spark mass heatmaps (section 1.8.), an exemplar correlative dataset from each of Ctrl-RV and MCT-RV was selected for this analysis. The rescaled and registered x and y coordinates of the centroid for each $Ca^{2+}$ spark was assigned a white pixel on an 8-bit array whose default pixel values were 0 and scaling was 5 nm/pixel. The arrays were read-in to a custom-written MATLAB programme which used the in-built routine that returns setblock values for quadtree decomposition (https://www.mathworks.com/help/images/ref/qtsetblk.html). A quad-tree map was generated from this which provided a minimum block size of 8×8 pixels. Regions of 8 or more for the smallest 4-connected quadrants, equivalent approximately to local density of $\geq 2$ sparks per 0.1 µm$^2$, were selected and the rest of the quadtree deleted. The resulting image represented the map of recurring spark sites (shown in Figure 3B-E and Suppl Fig S3). To interrogate their relationship to the local RyR organisation, the recurring spark site map was overlaid with the DNA-PAINT RyR image.

#### 1.10. Simulations

##### *1.10.1. Randomisation*

To interrogate the spatial relationship between the $Ca^{2+}$ spark positions and the nanoscale positions of RyR, two simulations were performed to include the spatial randomisation of these two variables in independent simulations using custom-written scripts implemented in IDL v 6.3 (Harris GeoSpatial). In the first, the x and y coordinates of $Ca^{2+}$ sparks in a correlative dataset acquired from Ctrl-RV myocytes were replaced with random x and y positions that encoded a different location of the cytoplasm of the cell. In each local sampling step, a spark footprint window was defined in the new, randomised position, but using the same width (i.e. equal to the FWHM) of the measured spark. In the latter randomisation, the total number of RyR puncta for each Ctrl-RV dataset was estimated and a new ensemble of RyR centroids, which matched the measured RyR number, was randomly positioned within the cell area (see Supplementary Figure S5).

##### *1.10.2. Resolution and Signal-to-noise ratio on alignment*

For simulating the impact of resolution and the signal-to-noise ratio (SNR), we simulated an averaged $Ca^{2+}$ spark image based on an experimentally recorded and rendered DNA-PAINT image of RyR on a rat ventricular myocyte. The simulated $Ca^{2+}$ spark image was randomly misaligned and then realigned using the alignment code. To investigate the impact of SNR on the alignment, the $Ca^{2+}$ image, traced from the DNA-PAINT image, was convolved by a 2D Gaussian function with a FWHM of 250 nm and then noise added such that the SNR would vary between 7.75 and 27.75 using the freely available ImageJ plugin, SNR, developed by Dr Daniel Sage ((8); http://bigwww.epfl.ch/sage/soft/snr/). The same plugin was used for estimating the SNR of the typical experimental, averaged $Ca^{2+}$ spark image (typically 15.5-18.0), as a reference point. In each of 10 iterations of re-alignment, the difference in each of the x and y coordinates was recorded as the alignment error and used for plotting Fig S4B. For simulating the impact of resolution, the simulated $Ca^{2+}$ image, traced from the DNA-PAINT image, was convolved with 2D Gaussian functions varying between 50 and 1000 nm. Noise was added to each simulated image to achieve an SNR of 17.0 before subjecting it to the same re-alignment iterations. Fig S4A show the violin plots of the alignment error in each of x and y dimensions after 10 iterations.

#### Supplementary Figures

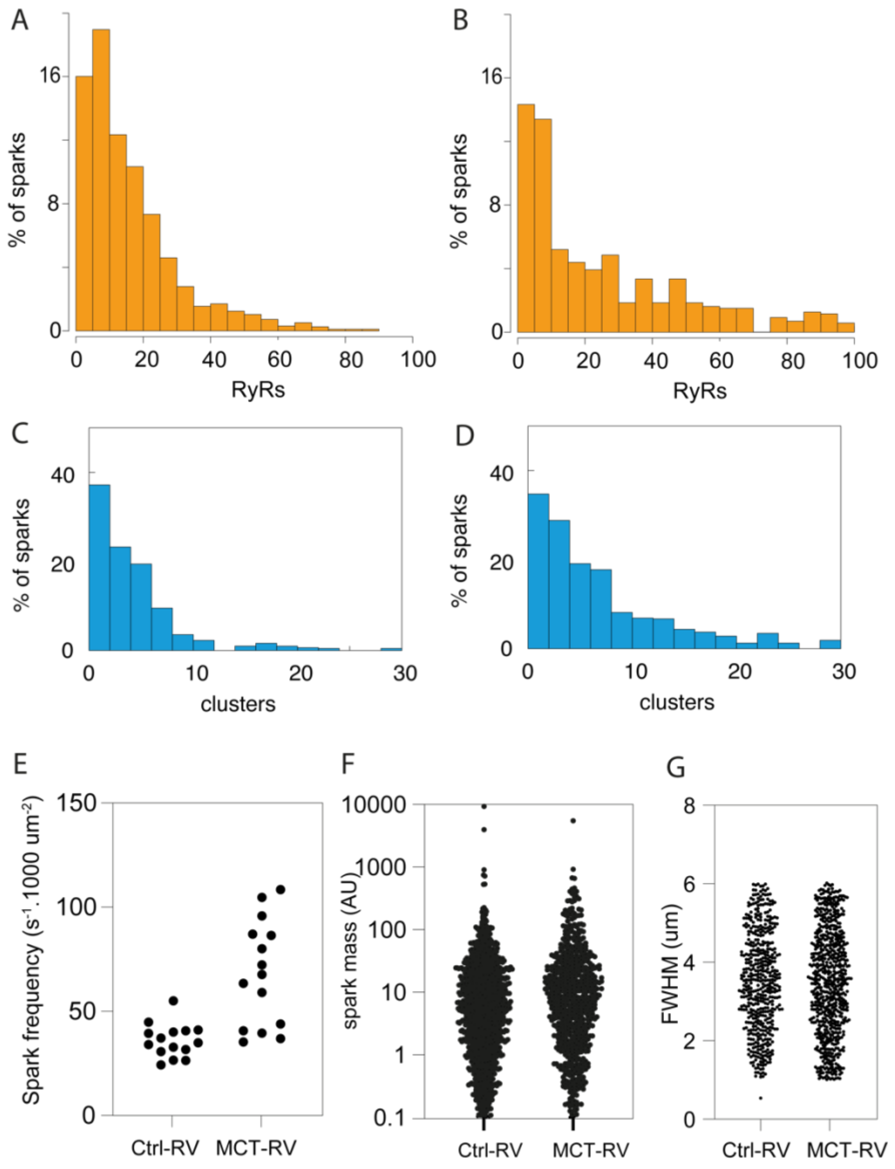


**Supplementary Figure S1. Statistics of local sampling of RyR organisation in the location of $Ca^{2+}$ sparks and spark properties in Ctrl-RV and MCT-RV myocytes. A&B.** Percentage histograms of the total RyR puncta count within the spark footprint in pooled Ctrl-RV (Mean ± SD: 18.1±16.4; median: 13, n=559 sparks; data pooled from 16 cells taken from 6 age-matched animals) and MCT-RV (23.5±17.7; median: 18, n=903 sparks, data pooled from 16 cells taken from 6 animals) datasets respectively. **C&D.** Percentage histograms of the total number of RyR clusters within the spark footprint in Ctrl-RV (5.16 ± 6.35; median: 3) and MCT-RV (8.11 ± 7.36; median: 5). **E.** A dot plot comparison of the mean spark frequency in Ctrl-RV (Median: 34.9 $s^{-1}$.1000 $\mu m^{-2}$; n =15 cells) and MCT-RV cells (median: 67.73 $s^{-1}$.1000 $\mu m^{-2}$; n=15 cells) analysed. Mann-Whitney test (p=0.0001, df=28). **F.** Dot plots of the spark mass (arbitrary units) of Ctrl-RV (Mean±SD 27.7±282.1) and MCT-RV (38.4±223.9; Mann-Whitney U test: p<0.0001; df=1992) sparks. G. FWHM of the Ctrl-RV (Mean: 3.50 µm; n=495 sparks) and MCT-RV (3.48 µm; n=684 sparks).

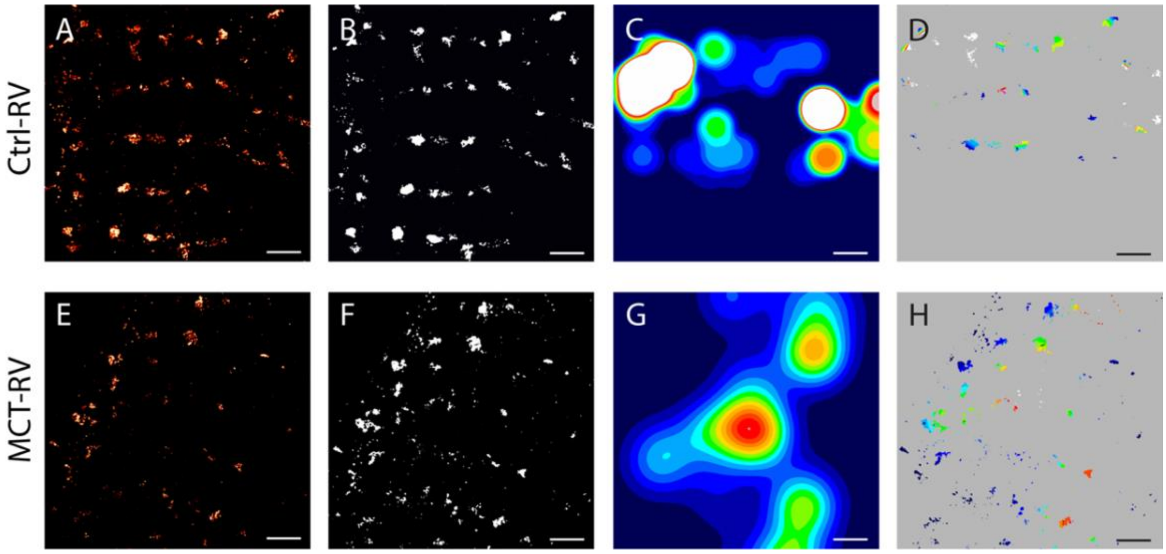


**Supplementary Figure S2. Generating the colour-coded DNA-PAINT RyR heatmaps encoding local average spark mass. A.** The greyscale-rendered DNA-PAINT image of sub-surface RyR pattern in a Ctrl-RV myocytes. **B.** A binary mask of the DNA-PAINT image. **C.** The 2D spark mass map **D.** The product of the RyR binary mask and the 2D spark mass map producing the average spark mass-coded RyR map for Ctrl-RV, illustrated in Fig 2 of the main manuscript. **E-H.** The equivalent pipeline for generating the spark mass-coded RyR heatmap for MCT-RV. Scale bars: 1 µm.

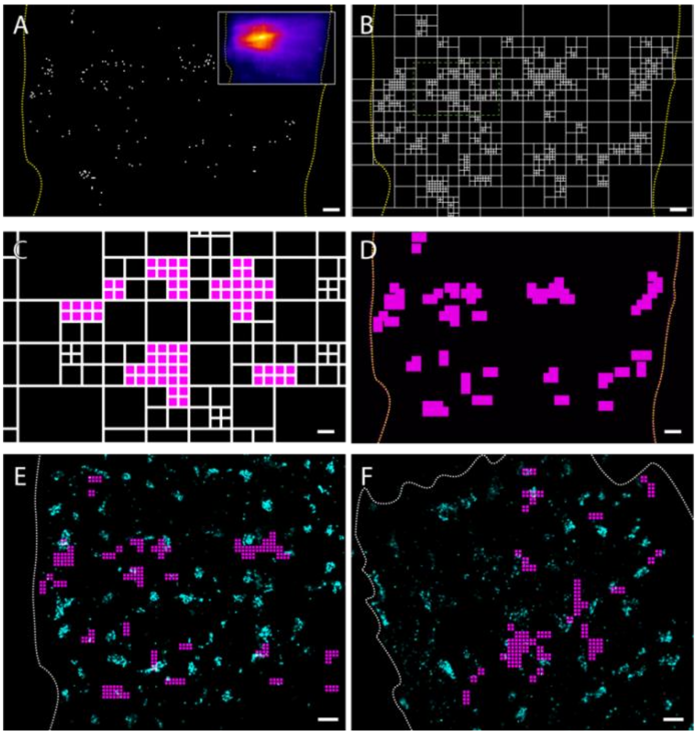


**Supplementary Figure S3. Quadtree visualisation of recurring spark sites. A.** A 2D pixel map of $Ca^{2+}$ sparks (example shown in inset) recorded over 20 s reveals a non-random pattern of spark locations. **B.** The quadtree decomposition created based on the pixel map. **C.** Selecting (coloured in magenta) ≥8 of the smallest 4-connected quadrants provide a local region that represents recurring spark sites. **D.** Shown is a zoomed-out view of the recurring spark sites across the full field of view.

**E&F.** Overlays of the recurring spark sites and the DNA-PAINT RyR images in Ctrl-RV and MCT-RV respectively. Scale bars: A,B,D: 1 µm, C: 60 nm, E&F: 250 nm

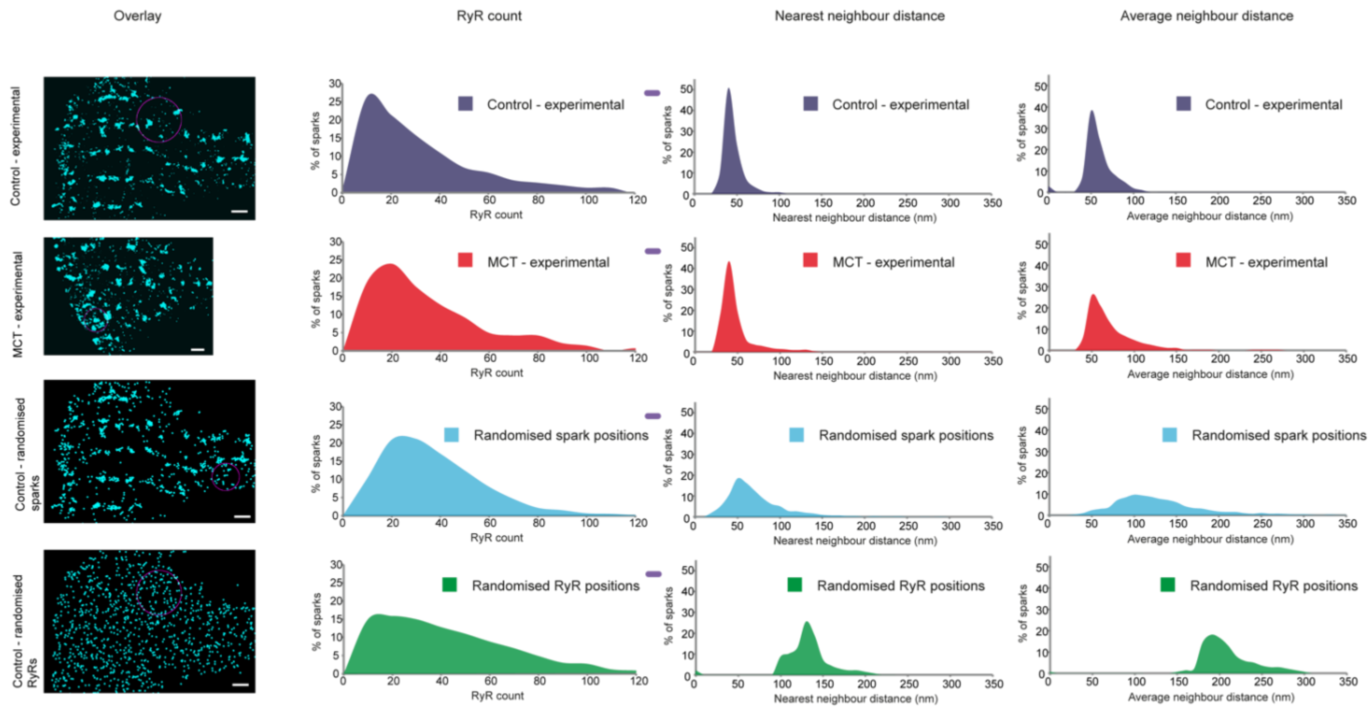


**Figure S4: Interrogating the relation between the $Ca^{2+}$ spark positions and sub-sarcolemmal RyR organisation through feature-randomisation.** Shown on the left column, are the four cases compared with overlays of either the RyR positions (cyan) overlaid with one of the $Ca^{2+}$ spark footprint windows (circle outlined by magenta line) used for sampling the RyR density or local organisation. The histogram shown in remaining columns, from left to right, show percentage histograms of the total RyR count, the mean nearest neighbour distance from each RyR, and the mean 3-neighbour average distance from each RyR respectively. The four scenarios considered in each row, from top to bottom, are the experimental spark and RyR positions in Ctrl-RV (n=169 sparks), the experimental spark and RyR positions in MCT-RV (n=263 sparks), Randomised spark positions interrogated by randomised spark positions in Ctrl-RV (n=1800 randomised sparks), and randomised spark and RyR positions based on Ctrl-RV cell geometry (n=1800 randomised sparks) respectively.

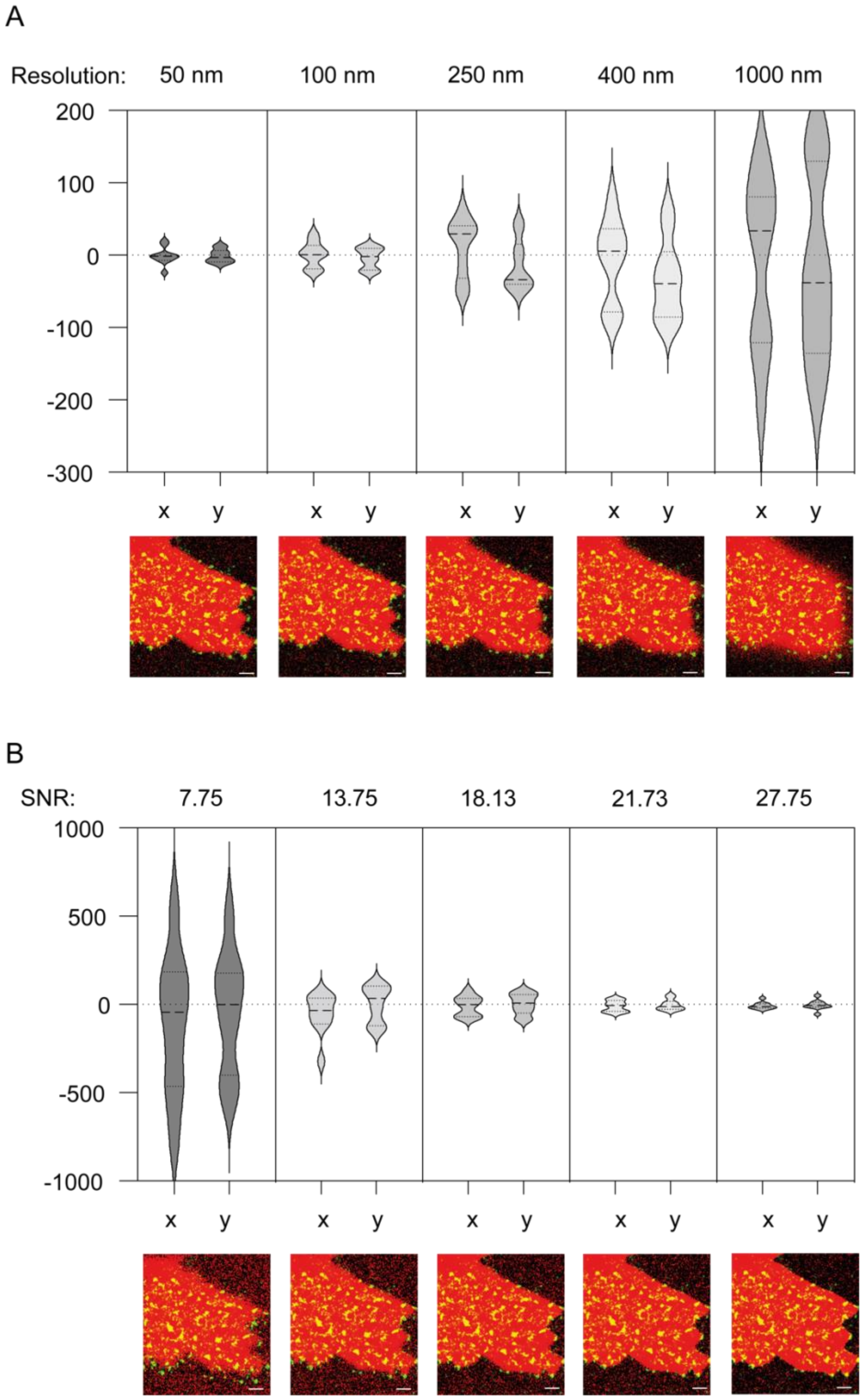


**Supplementary Figure S5: Simulation of the effect of resolution and signal-to-noise-ratio (SNR) on the alignment error. A.** Violin plots of the alignment error in the x and y dimensions in iterative alignment of a convolved copy (the FWHM of the convolution point spread function representing the 'Resolution' value indicated) of the simulated $Ca^{2+}$ image against the experimentally mapped RyR DNA-PAINT image. **B.** Violin plots of the alignment error in the x and y dimensions in iterative alignment of a noise-added copy of the simulated diffraction-limited $Ca^{2+}$ image against the experimentally mapped RyR DNA-PAINT image. Scale bar 1 µm. Each violin plot relates to 10 technical repeats of the simulation.

#### References


1. Fowler ED, Benoist D, Drinkhill MJ, Stones R, Helmes M, Wust RC, et al. Decreased creatine kinase is linked to diastolic dysfunction in rats with right heart failure induced by pulmonary artery hypertension. J Mol Cell Cardiol. 2015;86:1-8.

2. Jayasinghe I, Clowsley AH, de Langen O, Sali SS, Crossman DJ, Soeller C. Shining New Light on the Structural Determinants of Cardiac Couplon Function: Insights From Ten Years of Nanoscale Microscopy. Front Physiol. 2018;9:1472.

3. Steele EM, Steele DS. Automated detection and analysis of Ca(2+) sparks in x-y image stacks using a thresholding algorithm implemented within the open-source image analysis platform ImageJ. Biophysical journal. 2014;106(3):566-76.

4. Hurley ME, Sheard TMD, Norman R, Kirton HM, Shah SS, Pervolaraki E, et al. A correlative super-resolution protocol to visualise structural underpinnings of fast second-messenger signalling in primary cell types. Methods. 2021;193:27-37.

5. Marin Z, Graff M, Barentine AES, Soeller C, Chung KKH, Fuentes LA, et al. PYMEVisualize: an open-source tool for exploring 3D super-resolution data. Nat Methods. 2021;18(6):582-4.

6. Jungmann R, Avendano MS, Woehrstein JB, Dai M, Shih WM, Yin P. Multiplexed 3D cellular super-resolution imaging with DNA-PAINT and Exchange-PAINT. Nat Methods. 2014;11(3):313-8.

7. Baddeley D, Cannell MB, Soeller C. Visualization of localization microscopy data. Microsc Microanal. 2010;16(1):64-72.

8. Sage D, Unser M. Teaching image-processing programming in Java. IEEE Signal Processing Magazine. 2003;20(6):43-52.
